# The Gα_i/o_ GPCR LPAR2 Acts as a Non-Classical Factor for Aldosterone Induced Injury in a Cardiomyocyte Model

**DOI:** 10.64898/2026.09.29.755539

**Authors:** Saswat Kumar Mohanty, Rujun Gong, Marc Tatar

**Affiliations:** Department of Ecology, Evolution, and Organismal Biology, Center on the Biology of Aging, Brown University, Providence, RI 02912; Division of Nephrology, Department of Medicine University of Toledo College of Medicine, Toledo, OH 43606

**Keywords:** Aldosterone, LPAR2, GPCR, Mineralocorticoid receptor, AC16 human cardiomyocyte cells, Fibrosis

## Abstract

Aldosterone is a key regulator of systemic sodium, potassium, and fluid homeostasis. Chronic elevation of aldosterone, as observed in primary aldosteronism, chronic heart and kidney diseases, promotes cardiac hypertrophy, inflammation, fibrosis, and heart failure, which are further enhanced in aging. Although these pathological effects are primarily attributed to the mineralocorticoid receptor (MR), emerging evidence suggests roles for nonclassical G protein-coupled receptor (GPCR) signaling. We identified lysophosphatidic acid receptor 2 (LPAR2) as a novel GPCR candidate that mediates aldosterone signaling in cardiomyocytes. Using AC16 human cardiomyocyte cells and reporter gene assays, we demonstrate that aldosterone activates a Gαi/o-dependent GPCR pathway. Pharmacological inhibition of LPAR2 internalization confirms its specific reaction to aldosterone. Functional studies revealed that aldosterone-induced cardiomyocyte injury requires both MR and LPAR2, where specific antagonists to both receptors reduced the ability of aldosterone to induce collagen expression and cell death markers. These findings uncover a previously unrecognized non-classical mechanism of aldosterone action in the heart and highlight LPAR2 as a promising therapeutic target for aldosterone-driven cardiovascular disease.

## 1. Introduction

Aldosterone synthesized in the zona glomerulosa of the adrenal cortex is a regulator of systemic sodium, potassium, and fluid homeostasis when it activates through the mineralocorticoid receptor (MR) [1,2]. Chronic elevation of circulating aldosterone as observed in primary aldosteronism, chronic kidney disease, and heart failure is associated with increased risk of myocardial infarction, arrhythmia, and heart failure [3–5]. Aging may further contribute to these pathologies when aldosterone secretion becomes increasingly renin-independent, which elevates the aldosterone-to-renin ratio [6–8]. Prolonged aldosterone exposure promotes cardiomyocyte hypertrophy, oxidative stress, inflammation, and deposition of extracellular matrix. These events contribute to myocardial fibrosis and impair heart function. [9].

The classic action of aldosterone is mediated by nuclear MR [1,10,11]. Ligand bound MR translocates to the nucleus to regulate the transcription of genes involved in inflammation, transport, and tissue remodeling [1]. Accordingly, in many conditions, pharmacological inhibition of MR by antagonist such as spironolactone and eplerenone significantly reduce cardiovascular morbidity and mortality, yet there remains a wide appreciation that additional aldosterone responsive signaling contributes to cardiac pathology even when MR is inhibited [12–14]. Numerous studies suggest that aldosterone has non-genomic action which elicits a rapid cellular response. These responses include the activation of PI3K/Akt, protein kinase C, ERK1/2, intracellular calcium mobilization, ROS production, and modulation of cAMP signaling [10,15]. This rapid response has been reported in endothelial cells, renal epithelial cells, vascular smooth muscle cells, and cardiomyocytes, and suggest there is some form of membrane associated aldosterone signaling [15–17].

Despite more than two decades of investigation, the identity and function of a membrane receptor system responsible for these rapid actions remains controversial. Several candidates have been proposed, including G protein-coupled estrogen receptor (GPER), angiotensin II type 1 receptor transactivation, epidermal growth factor receptor signaling, and membrane-associated MR complexes [18–20]. However, the receptor and signaling mechanisms underlying the rapid aldosterone actions have been difficult to establish and appear to vary across tissues and cell types. There may be no single mechanism to fully account for the spectrum of aldosterone-induced rapid signaling [21–23]. The versatility of aldosterone was even apparent in a laboratory study of the fruit fly *Drosophila melanogaster*. We previously found that aldosterone induces cardiorenal fibrosis in adult Drosophila through action that requires the *Drosophila* GPCR dopamine ecdysone receptor (dopEcR) expressed in fly cardiomyocytes [24]. This observation motivates our current question: identify GPCR candidate(s) that mediate aldosterone-induced cardiac fibrosis in the mammalian heart.

G protein-coupled receptors (GPCRs) are a large family of cell-surface signaling receptors that modulate virtually every aspect of cardiovascular physiology and disease [25]. GPCRs transduce extracellular signals by coupling to heterotrimeric G proteins containing Gα subunits from four principal families—Gα_s_, Gα_i/o_, Gα_q/11_, and Gα_12/13_, which engage distinct downstream effector pathways [26]. Activated Gα_i/o_-coupled GPCRs suppress adenylyl cyclase activity, decrease intracellular cyclic AMP levels, and activate downstream pathways involved in cell proliferation, survival, migration, and extracellular matrix remodeling [27]. Lysophosphatidic acid receptor 2 (LPAR2) is a member of the endothelial differentiation gene (EDG) family of lysophosphatidic acid receptors, which includes LPAR 1-3. LPAR2 is a ‘Class A’ GPCR activated by bioactive phospholipid lysophosphatidic acid (LPA). This receptor can couple to Gα_i/o_, in addition to Gα_q/11_ and Gα_12/13_ [28]. LPAR2 signaling promotes cell survival, cytoskeletal reorganization, inflammatory responses, and fibrotic remodeling by activating multiple downstream pathways, including RhoA/ROCK, MAPK, PI3K/Akt, and serum response factor signaling. LPA-LPAR signaling is implicated in cardiac fibrosis, vascular remodeling, and myocardial injury [28–30]. Here we find that LPAR2 participates in aldosterone signaling, alongside its conical activity induced by LPA. Our work provides a new example for how steroid hormone signaling and GPCR-mediated pathways converge to regulate cardiovascular function. Such systems may work via crosstalk between nuclear receptors and GPCRs to integrate rapid membrane-initiated signaling with slower transcriptional responses [18,31].

Initially we identified LPAR2 as a candidate for aldosterone responsive GPCRs in human embryonic kidney cells (HEK293). We then studied AC16 human cardiomyocytes to understand impacts of aldosterone that require LPAR2. AC16 cells are primary adult human ventricular cardiomyocytes hybridized with SV40-transformed human skin fibroblasts. AC16 cells retain some of the genetic traits of fibroblast [32]. We used reporter gene assays, cyclic AMP measurements, receptor internalization analyses, and functional studies in these cells to demonstrate that aldosterone activates LPAR2-dependent Gα_i/o_ signaling. We show that although LPAR2 mediates rapid non-genomic signaling in response to aldosterone, both LPAR2 and MR are required for aldosterone to induce cardiomyocyte injury, fibrosis, and hypertrophy. These observations establish a previously unrecognized LPAR2-MR signaling axis and provide potential insights into aldosterone-induced cardiovascular remodeling.

## 2. Materials and methods

### 2.1. Cell culture maintenance and cardiomyocyte differentiation

Human embryonic kidney (HEK293) cells and AC16 human cardiomyocyte cells were obtained from the American Type Culture Collection (ATCC). ΔGi HEK cells were donated by Prof. Asuka Inoue, from Graduate School of Pharmaceutical Science, Tohoku University [33]. HEK293 and ΔGi HEK cells were maintained in Dulbecco’s Eagle’s Medium (DMEM, high glucose, GlutaMAX™ Supplement, pyruvate; Gibco cat no. 10569010) supplemented with 10% fetal bovine serum (FBS) and 1% antibiotic-antimycotic solution (Gibco 15240062) in a humidified atmosphere of 5% CO_2_ at 37°C. Similarly, AC16 cells were maintained in DMEM/F-12 media supplemented with 12% FBS and 1% antibiotic-antimycotic solution with 5% CO_2_ at 37°C. To induce cardiomyocyte-like phenotypes from AC16 cells, cultures were transferred to differentiation medium supplemented with 1% horse serum, 1X ITS, 10nM all trans retinoic acid (ATRA) and 1% antibiotic-antimycotic solution in DMEM/F-12 for 4 days. These differentiated cells were used to study downstream pathological, fibrotic and gene/protein expression. Prior to all experimental treatments, cells were first serum-starved in opti-MEM medium for 12-16 h to eliminate baseline growth factor signaling.

### 2.2. GPCR reporter gene expression assay

We used the transcriptional reporter gene expression Promega Luciferase Assay System to identify the activation of GPCR following aldosterone stimulation in HEK293 cells. Cells were transfected with reporter vectors using TransIT-LT1 transfection reagent for 24 h in a 96-well plate. Vector details are available in supplementary materials. Transfected cells were stimulated by aldosterone (or antagonist) for each experimental time point. In cases where we first inhibit a receptor, specific antagonists were applied to cells 30 min prior to aldosterone stimulation. Reporter firefly luciferase activity and control nanoluc luciferase (Thymine Kinase) were measured using the Promega Nano-Glo® Dual-Luciferase® Reporter Assay System on a Spectramax M5 multi-mode microplate reader. Data were expressed as luminescence unit (RLU) relative to control vector.

### 2.3. Intracellular cAMP accumulation assay

Intracellular cyclic adenosine monophosphate (cAMP) accumulation was quantified using the Promega cAMP-Glo™ Assay. Cells were stimulated with adenylyl cyclase activator forskolin (10 μM) [34,35] within each experimental treatment. Cells treated with forskolin alone were used as positive controls to quantify maximum cAMP induction. Luminescence was measured using the Spectramax multimode luminometer. To account for assay variability and establish physiological scaling, final data were calculated and expressed as a percentage of the maximum cAMP accumulation stimulated by forskolin [36,37].

### 2.4. Bioluminescent LPAR2 receptor internalization assay

LPAR2 cell surface expression and internalization kinetics were quantified using the Promega Nano-Glo® HiBiT Extracellular Detection System. To our specifications, HiBiT-tagged human LPAR2 vector were produced by Promega, USA **(Supplementary materials)**. AC16 human cardiomyocyte cells expressing HiBiT-tagged human LPAR2 were treated with aldosterone and/or LPAR2 specific agonist (GRI977143) in the presence or absence of antagonist for each experimental time point. At each time point, cell culture medium was replaced with Nano-Glo® HiBiT Extracellular Detection Reagent containing the membrane-impermeable LgBiT protein and Nano-Glo® substrate. The measured luminescence is proportional to the extracellular expression of LPAR2. A decrease in extracellular luminescence represents internalization of the receptor. Receptor internalization was calculated as percent change in extracellular RLU.

### 2.5. Transient overexpression of human LPAR2

LPAR2 was overexpressed for functional assays from a sequence-verified Human Tagged ORF Clone encoding EDG4/LPAR2 (NCBI Reference Sequence: NM_004720) from OriGene Technologies, Inc **(Supplementary materials)**. AC16 cells were expanded to 70–80% confluence before transfection. OriGene LPAR2 plasmid or an empty vector control (pCMV6 backbone) was transiently delivered using TransIT-LT1 transfection reagent at a ratio of 3:1. Cells were incubated with the transfection complexes for 24–48 hours to ensure robust expression of the target receptor prior to assays.

### 2.6. Western Blotting for Injury and Fibrosis Markers

Differentiated AC16 cells were lysed in ice-cold RIPA lysis buffer supplemented with protease and phosphatase inhibitors. Protein (25 μg) was separated in 4-15 % tris-glycine gel and transferred to immobilon-P PVDF membrane (Roach, USA). After blocking with 5% nonfat skimmed milk for 1 h at room temperature, membranes were incubated with primary antibodies targeted for Collagen Type I (Collagen 1), Cardiac Troponin I (cTnI), Natriuretic Peptide A (NPPA) and α-Tubulin at 4 °C overnight. Membranes were subsequently washed and incubated with fluorescent tag secondary antibodies. Protein bands were visualized using the Bio-Rad chemidoc MP detection system. Band densities were quantified using ImageJ software and normalized to α-Tubulin as loading control.

### 2.7. Immunocytochemistry / Immunofluorescence (ICC/IF) for Collagen 1

To visualize intracellular Collagen 1 expression, AC16 cells were seeded on Gelatin coated sterile coverslips and differentiated for 4 days. Cells were stimulated with aldosterone (1 μM) for 24 h in the presence or absence of LPAR2 antagonist (LPA2 antagonist1, 30 nM) or MR antagonist (Eplerenone, 5 μM). Treated cells were fixed with 4% paraformaldehyde for 10 min at room temperature. After washing with PBS, cells were permeabilized with 0.1% Triton X-100 for 15 min and blocked with 5% BSA in PBS for 1 h. Coverslips were then incubated with primary antibody targeting Collagen 1 at 4°C overnight. The cells were washed and incubated with Alexa 488 conjugated secondary antibody and phalloidin for 1 h at room temperature. The coverslips were mounted onto glass slides using VECTASHIELD HardSet Antifade Mounting Medium with DAPI. The images were captured using Zeiss LSM 880 Confocal Microscope, and the corrected total cell fluorescence (CTCF) of collagen 1 was quantified using ImageJ software and normalized to number of nuclei.

### 2.8. Real-Time Quantitative PCR (RT-qPCR)

Total RNA was isolated from cells using the TRIzol reagent method via chloroform phase separation and isopropanol precipitation [38]. To ensure complete elimination of potential genomic DNA contamination, the isolated total RNA was subjected to post-isolation digestion using the TURBO DNA-free™ Kit (Invitrogen/Thermo Fisher Scientific). Purified RNA was reverse transcribed into cDNA using the Bio-Rad iScript™ cDNA Synthesis Kit. cDNA was subjected to qPCR using SYBR green master mix using an Applied Biosystems QuantStudio 5 Real-Time PCR System for 40 cycles. mRNA was quantified via the 2^−ΔΔCt^ method, with GAPDH as an internal control [39]. Primer details are listed in supplementary materials.

### 2.9. LDH Leakage Cytotoxicity Assay

Lactate dehydrogenase (LDH) leakage into the culture supernatant was measured to quantify cardiomyocyte plasma membrane rupture using the Thermo Scientific Pierce™ LDH Cytotoxicity Assay Kit. Cell culture supernatant (50 μL) from each treatment well was transferred to a fresh 96-wel plate and mixed with 50 μL of LDH reaction mixture. The plate was incubated at room temperature for 10 min protected from light. The reaction was stopped by adding 50 μL stop solution. Fluorescence was measured using excitation of 560 nm and emission of 590 nm. Cytotoxicity was estimated asTthe ratio of spontaneous LDH release to maximum LDH release.

### 2.10. Multiplexed ATP Cell Viability and CellTox™ Green Cytotoxicity Assays

A multiplexed protocol combining the CellTox™ Green Cytotoxicity Assay and the CellTiter-Glo® Luminescent Cell Viability Assay (Promega Corporation, Madison, WI, USA; Catalog Nos. G8741 and G7570) simultaneously monitors cell viability and plasma membrane degradation in the same experimental well.

### 2.11. Cellular Reactive Oxygen Species (ROS) Detection Assay

Intracellular reactive oxygen species (ROS) production was monitored fluorometrically using an H2DCFDA - Cellular ROS Assay Kit (Abcam, Cambridge, UK; Catalog No. ab113851).

### 2.12. Statistical Analysis

All data are expressed as mean ± SEM from at least 3 independent biological replicates (n ≥ 3). Statistical analysis between multiple groups were evaluated using one-way and two-way analysis of variance (ANOVA) as applicable. A p-value ≤ 0.05 was considered statistically significant. All statistical calculations, charts, and data plots were generated using GraphPad Prism software (GraphPad Software, Boston, MA, USA).

## 3. Results

### 3.1. Aldosterone activates Gαi/o coupled GPCR (LPAR2) in HEK293 cells

HEK293 cells were used to initially identify GPCRs responsive to aldosterone. In cells treated with aldosterone, we screened reporter vectors specific to GPCR signaling pathways [40]. Following aldosterone exposure at varying concentrations and time points, we measured firefly luciferase expression modulated by transcription factor response elements uniquely associated with Gα_s_, Gα_i/o_, Gα_q/11_, and Gα_12/13_ signaling. Aldosterone selectively induced luciferase expression driven by the SRE response element in a dose and time dependent manner with a maximum activity at 10^-6^ M (1 μM) for 6 h treatment (**Fig. 1C**). Because SRE is activated by Gα_i/o_, we infer that Aldosterone activates downstream Gα_i/o_ coupled GPCR signaling [40]. To validate this Gα_i/o_ coupled GPCR activated signaling we measured how aldosterone, in a dose dependent manner after 6 h, inhibits the accumulation of cAMP. Aldosterone at 10^-6^ M A significantly reduces cAMP, indicative of activated Gα_i/o_ (**Fig. 1E**). We further show that aldosterone requires Gα_i/o_ in tests with cells that lack this Gi, ΔGi HEK cells [33]aldosterone could not inhibit cAMP (**Fig. 1F**). The next goal was to identify which Gα_i/o_ coupled GPCR activates SRE in HEK cells. HEK293 cells express five functional Gα_i/o_ dependent GPCRs: A2B adenosine receptor (A2B), Lysophosphatidic acid receptor 1 & 2 (LPAR 1 & 2), Somatostatin receptor 5 (SSTR5) and Protease-activated receptor 1(PAR1) [41]. We rule out PAR1 because it is activated upon the proteolytic cleavage at its C-terminus end by its ligand, a function lacking in aldosterone [42]. To test the other candidates of SRE activation, we treated HEK 293 cells with varying concentrations of receptor-specific antagonists (alone) and measured SRE-driven luciferase activity. Only receptor specific inhibition of LPAR1 and LPAR2 significantly reduce the SRE activity **(Supplementary Data)**. To evaluate the impact of aldosterone on SRE through these LPARs, cells were pretreated for 30 minutes with IC50 concentrations of either LPAR1 or LPAR2 antagonists prior to aldosterone exposure (10^-6^ M). SRE-luciferase activity and cAMP accumulation were quantified after 6 hours. Uniquely, pharmacological inhibition of LPAR2 abolished aldosterone-induced SRE activation and cAMP suppression, suggesting that aldosterone acts via the Gα_i/o_ coupled receptor LPAR2 (**Fig. 1G-I**).

**Figure 1:**
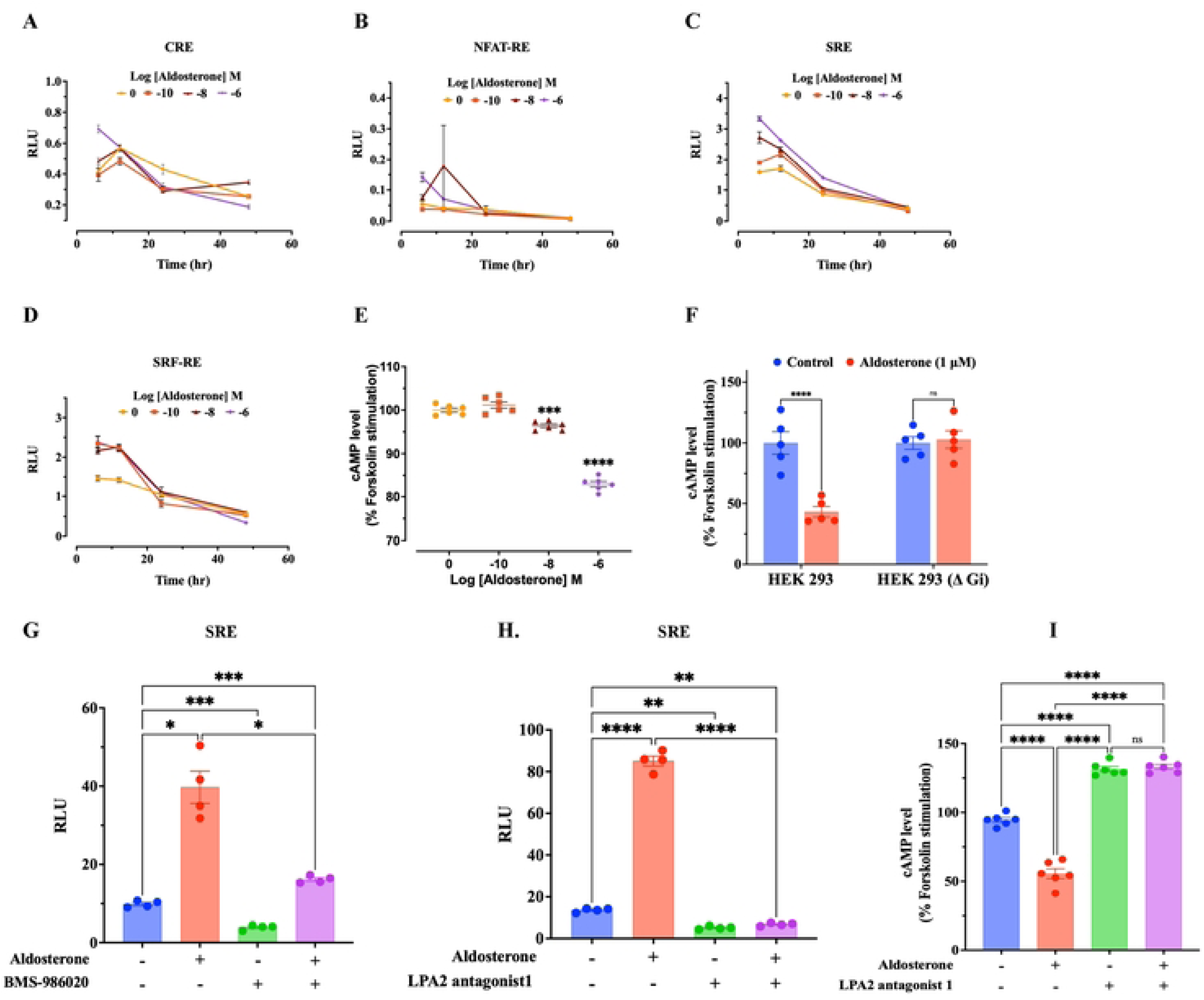
Identification of the GPCR candidate activated by Aldosterone stimulation HEK293 cells. Aldosterone induce SRE reporter gene activity in a dose and time dependent manner and significantly inhibited forskolin-stimulated cAMP accumulation indicating activation of G_αi_ coupled GPCR. Further, pharmacological inhibition of receptors by specific antagonist identified the involvement of LPAR2 GPCR. **A-D:** Reporter gene screening as function of time and aldosterone stimulation, **E:** measurement of cAMP accumulation after 6 h of increasing concentration of aldosterone stimulation, **F:** measurement of cAMP accumulation after 6 h of aldosterone (1μM) stimulation in wild type and KO HEK293 cells, **G-H:** SRE reporter gene expression in cells treated with aldosterone (1μM) in presence or absence of BMS-986020 (LPARI antagonist) and LPA2 antagonist 1, **I:** cAMP accumulation level in aldosterone (1μM) stimulated cells in presence or absence of LPA2 antagonist 1. Asterisks *, **, ***, **** indicates the significance level at *p* ≤0.05, *p* ≤0.01, *p* ≤ 0.001, *p* ≤0.0001, respectively.

### 3.2. Aldosterone induces LPAR2 activation and internalization in AC16 human cardiomyocytes

We explored LPAR2 in AC16 human cardiomyocyte cells to study potential novel impacts of aldosterone on the heart. SRE was increased and cAMP was suppressed when AC16 cells were treated with aldosterone (**Fig. 2A, B**). In contrast to the relatively fast response observed in HEK 293 cells, the effect of aldosterone on SRE and cAMP in AC16 cells required 12 h treatment at 10^−6^M (1μM) and was not seen at earlier time points. SRE and cAMP were not modulated by aldosterone if cells were first treated with an IC50 concentration of LPAR2 antagonist **(Supplementary Data)** prior to the aldosterone treatment (**Fig. 2C, D**). To further investigated the connection between aldosterone and LPAR2 in AC16 cells, we followed LPAR2 internalization, which is characteristic of agonist activated GPCRs [43,44]. We evaluated receptor endocytosis of a transfected HiBiT-LPAR2 fusion vector (Promega). As a positive control, the LPAR2-specific agonist GRI977143 demonstrated time- and dose-dependent receptor internalization at 30 min and 1 h, and at an EC50 of 1μM (**Fig. 2E**). Subsequently, HiBiT-LPAR2 expressing AC16 cells were treated with 1μM aldosterone, GRI977143 or both in combination for 1h, and in each case in the presence or absence of LPAR2 antagonist. Aldosterone alone did not induce internalization, whereas LPAR2 agonist GRI977143 alone induced 50% internalization after 1h. Importantly, receptor internalization was strongly increased when LPAR2 agonist was combined with aldosterone. There is synergy between these factors. Conversely, cells treated with the LPAR2 antagonist showed less internalized receptor in response to GRI977143, and less synergy between aldosterone and GRI977143 (**Fig. 2F**).

**Figure 2:**
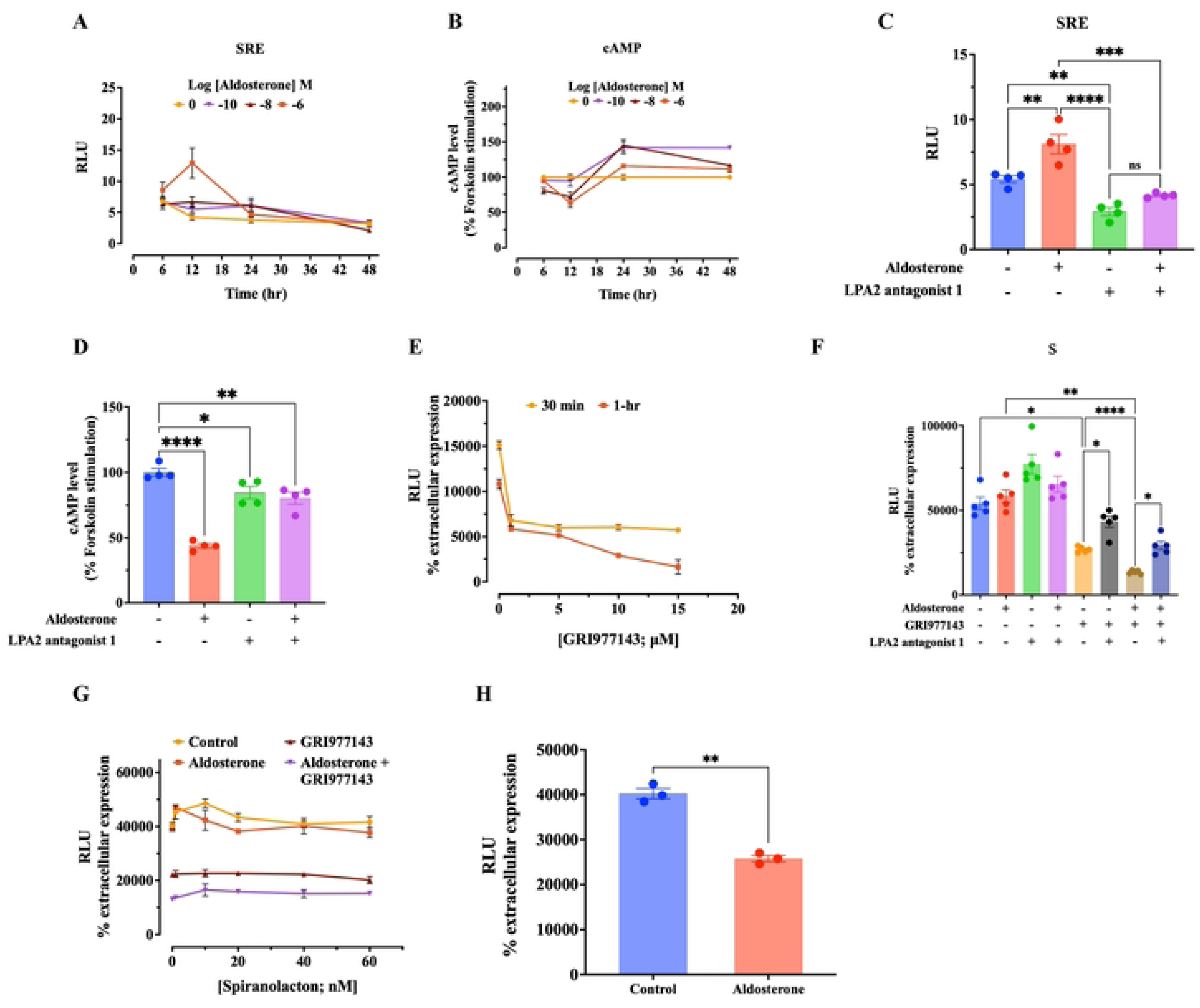
Validation of the activation of G_αi_ coupled GPCR, LPAR2 activation by aldosterone in AC16 human cardiomyocyte cells. Aldosterone induced SRE and reduction in cAMP accumulation was observed after 12 h at only 1μM concentration. Pharmacological inhibition by LPA2 antagonist 1 blocks the effect of aldosterone. Aldosterone induces LPAR2 receptor internalization after 2 h, however synergistically it showed additive efect with LPAR2 agonist (GRI97143). The internalization was not inhibited by MR antagonist spironolactone. **A-B:** Effect of aldosterone on SRE and cAMP as a function of time and dose respectively, **C-D:** Effect of aldosterone (1μM) on SRE and cAMP respectively, in presence or absence of LPA2 antagonist 1, **E:** Dose and time dependent effect of GRI977I43 on LPAR2 internalization, **F:** LPAR2 internalization after Ih, **G:** LPAR2 internalization in presence or absence of spironolactone after 1 h, **H:** LPAR2 internalization upon aldosterone(1μM) for 2 h. Asterisks *,**,***,**** indicates the significance level at *p* ≤0.05, *p* ≤0.01, *p* ≤0.001, *p* ≤0.0001, respectively.

It is important to also consider how the mineralocorticoid receptor (MR) may mediate the LPAR2 internalization induced by agonist and aldosterone. We pretreated the AC16 cells with varying concentrations of the MR antagonist spironolactone prior to testing how aldosterone and GRI977143 impact LPAR2 internalization. Despite the MR inhibition, LPAR2 was internalized at 1 h to the expected level induced by the combined exposure of GRI977143 and aldosterone (**Fig. 2G**). Unexpectedly, aldosterone alone in these trials induced LPAR2 internalization following a 2 h treatment, unlike the previously tested 1 h timepoint (**Fig. 2h**).

### 3.3. Synergy of aldosterone and LPAR2 agonist GRI977143

Following the observed interaction between aldosterone and GRI977143 upon receptor internalization, we evaluated their combined effect on SRE reporter activity and cAMP levels. We hypothesized that co-treatment of AC16 cells with GRI977143 and aldosterone would accelerate how aldosterone alone modulates SRE (increased) and cAMP (reduced) pathways after 12 h. We treated AC16 cells with aldosterone, GRI977143, or their combination in the presence or absence of the LPAR2 antagonist for 3 and 6 h. GRI977143 alone elicited a significant signaling response; aldosterone monotherapy failed to produce any detectable effect; co-treatment of agonist and aldosterone significantly accelerated the response compared to GRI977143 treatment alone. All responses were blocked upon pretreatment with LPAR2 antagonist (**Fig. 3A**).

**Figure 3:**
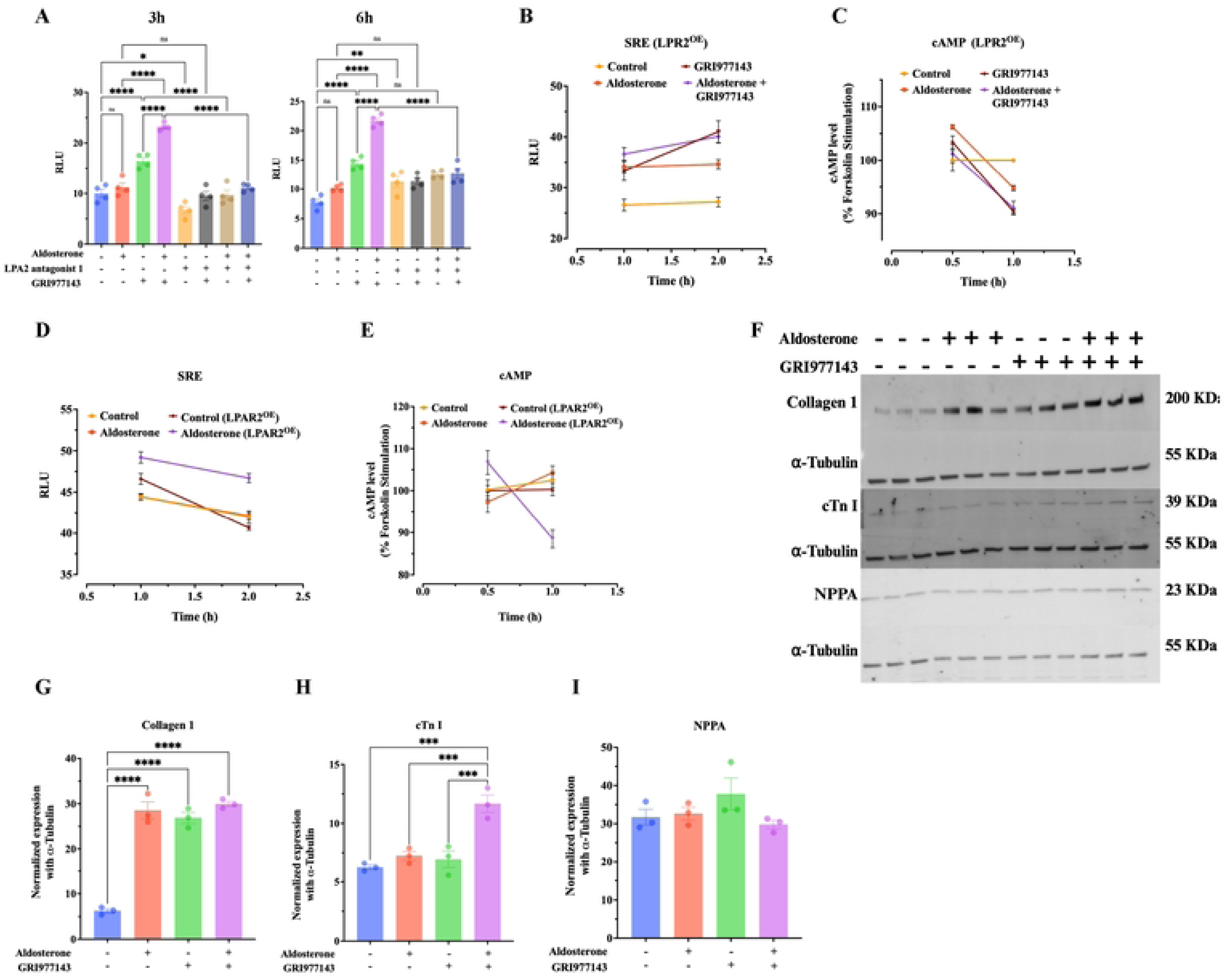
The cell surface expression level of LPAR2 is responsible for aldosterone mediated signaling in AC16 human cardiomyocyte cells. Aldosterone exerts its effect on LPAR2 more efficiently when the cells express high LPAR2 than the normal basal level expression. This shifts the effect of aldosterone from 12 h to 1 h. As differentiated AC 16 cells express more LPAR2 than the proliferating cells, we tested its effect on fibrotic protein like collagen. Differentiated AC 16 cells express significantly high level of collagen-1 protein after 24 h of aldosterone (1μM) stimulation. A synergistic effect of aldosterone with GRI977143 was observed for cTN I protein expression. **A:** Synergistic effect of aldosterone and GRI977143 after 3 h and 6 h of stimulation in proliferative AC 16 cells, **B, C:** SRE reporter and cAMP level in cells overexpressed with LPAR2 after single treatment with aldosterone and GRI977143 or combined treatment respectively, **D, E:** Effect of aldosterone on SRE and cAMP in wild type and LPAR2 overexpressed cells, F: Western blot representation of injury and fibrotic markers in differentiated AC 16 human cardiomyocyte cells, **G-I:** Densitometric analysis of western blot protein expression of collagen 1, cTn I, and NPPA respectively. Asterisks *,**,***,**** indicates the significance level at*p* ≤0.05, *p* ≤0.01, *p* ≤0.001, *p* ≤0.0001, respectively.

To further evaluate the impact of aldosterone via LPAR2 we transiently overexpressed LPAR2 in AC16 cells. In these cells, aldosterone and GRI977143 equally increased SRE reporter activity after 1 h. SRE reporter activity plateaued at 2 h in aldosterone-treated cells, whereas it was significantly higher in GRI977143-treated cells. No significant reduction in cAMP was observed at 1 h with aldosterone or GRI977143, but both induced a strong decrease after 2 h. In AC16 cells overexpressing LPAR2, co-treatment with aldosterone and GRI977143 altered SRE activity and reduced cAMP at 2 h to levels seen with GRI977143 alone (**Fig. 3B, C**). Crucially, LPAR2-overexpressing cells treated with aldosterone alone induced SRE reporter activity and cAMP levels to a greater extent than aldosterone treatment of wild-type (non-overexpressing) cells (**Fig. 3C, D**). Overall, these findings suggest that the expression of exogenous LPAR2 facilitates aldosterone-induced downstream signaling.

### 3.4. Aldosterone Induces Cardiomyocyte pathologies through LPAR2 and the mineralocorticoid receptor

To understand potential physiological impacts of aldosterone via LPAR2, we investigated pro-fibrotic and hypertrophy markers (Collagen 1, cTnI, and NPPA) in differentiated AC16 cardiomyocytes, which express approximately six-fold higher levels of LPAR2 mRNA relative to their undifferentiated counterparts **(Supplementary Data)**. In these differentiated cells, aldosterone and GRI977143 alone significantly upregulated collagen 1 protein when treated for 24 hours (**Fig. 3F**), while combined treatment did not further increase expression. In contrast, we see a clear synergistic effect in cTnI protein expression, because no single treatment elevated the protein, but combined treatment induced a marked increase. No treatment affected the level of NPPA protein (**Fig. 3G-I**). This indicates that aldosterone is sufficient to enhance a fibrotic marker (collagen-1), while the LPAR2 agonist may catalyze the aldosterone response.

We measured additional markers of cell fitness including cell survival, an index of cardiac hypertrophy and collogen accumulation.

A 24 h aldosterone treatment significantly reduced differentiated AC16 cell survival, decreasing viability by 16.8% relative to untreated control. Treatment with the GRI977143 alone, and when combined with aldosterone, induced a comparable decrease in cell viability (**Table 1**). Direct LPAR2 stimulation copies the cytotoxic response induced by aldosterone alone. Cardiomyocyte injury also associates with cardiac hypertrophy, characterized by increased intracellular MHCβ (MYH7) and reduced MHCα (MYH6), which contributes to myocardial stiffness in the human heart [45]. To evaluate this hypertrophic response, we measured MYH7 and MYH6 mRNA in differentiated AC16 cells following 24 h of aldosterone exposure. Aldosterone treatment significantly increased MYH7 mRNA while leaving MYH6 mRNA expression unchanged. Aldosterone increased the MYH7:MYH6 expression ratio (hypertrophic index) (**Fig. 4A, B**), consistent with cardiac hypertrophy when evaluated in clinical settings [45]. Notably, pharmacological inhibition with an LPAR2 antagonist alone or the MR antagonist eplerenone prevented the upregulation of MYH7 by aldosterone and restored the MYH7:MYH6 ratio toward baseline (**Table 2**).

**Figure 4:**
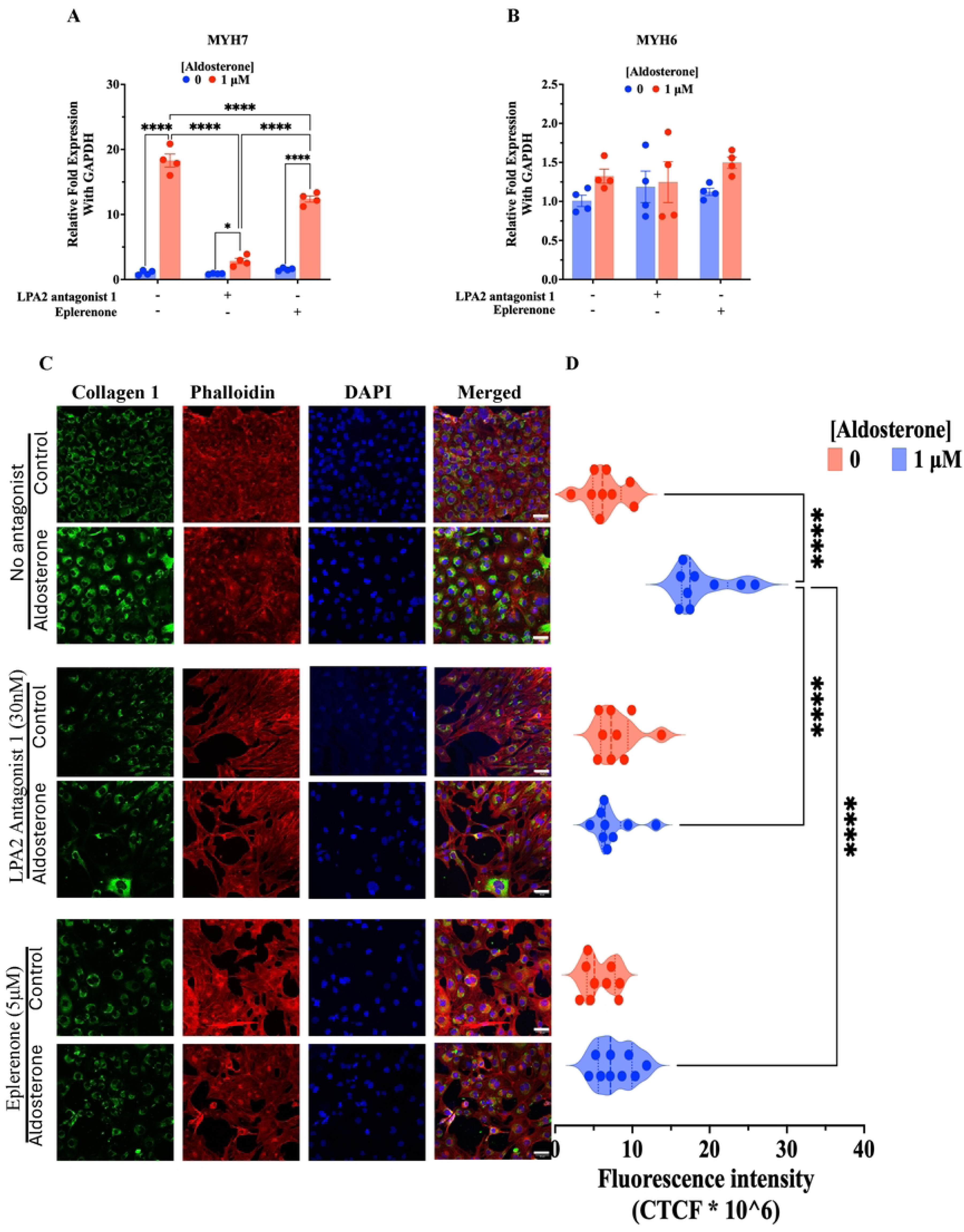
Both MR and LPAR2 are responsible for Aldosterone-Induced Cardiomyocyte Hypertrophy and Pro-Fibrotic Signaling in differentiated AC16 human cardiomyocyte cells. Aldosterone (IμM) induces cardiomyocyte hypertrophy as evidenced from the increased mRNA expression of MYH7 (β-MHC). Moreover, it also increases intracellular collagen 1 expression. All these effects were abolished by pharmacological inhibition of either MR or LPAR2 antagonist. **A & B:** mRNA expression of MYH7 and MYH6 respectively, **C:** ICC/IF image of collagen 1 in differentiated AC 16 cells, **D:** Densitometric graph for fluorescent intensity. Asterisks **♦* indicates the significance level at *p* ≤ 0.0001.

We further examined the pro-fibrotic state of differentiated AC16 response by measuring intracellular collagen 1 accumulation using immunocytochemistry. Cells were pretreated with either vehicle, LPAR2 antagonist (30 nM) or eplerenone (5 μM) for 30 min, then treated with aldosterone for 24 h. Aldosterone induced a significant increase in intracellular collagen 1 protein, while pretreatment with either LPAR2 antagonist or MR antagonist blocked this increase (**Fig. 4C, D**).

Cytotoxicity results in plasma membrane integrity disruption, which can be measured by lactate dehydrogenase (LDH) leaked into the culture medium. Extracellular LDH levels in differentiated AC16 cells were markedly elevated from cells treated with aldosterone for 24 h. To assess the relative roles of MR and LPAR2, cells were pretreated with either the LPAR2 antagonist or the MR antagonist eplerenone (respectably at 1 μM and 5 μM). Both inhibitors attenuated the release of LDH induced by aldosterone. Notably, pretreatment with LPAR2 antagonist completely blocks the LDH leakage, demonstrating that this GPCR is required for aldosterone-induced membrane injury (**Fig. 5B**). AC16 cells were also treated with the LPAR2 agonist GRI977143 as a positive control. As expected, this elevated LDH release, and the impact was fully blocked by co-treatment with LPAR2 antagonist (**Fig. 5A**).

**Figure 5:**
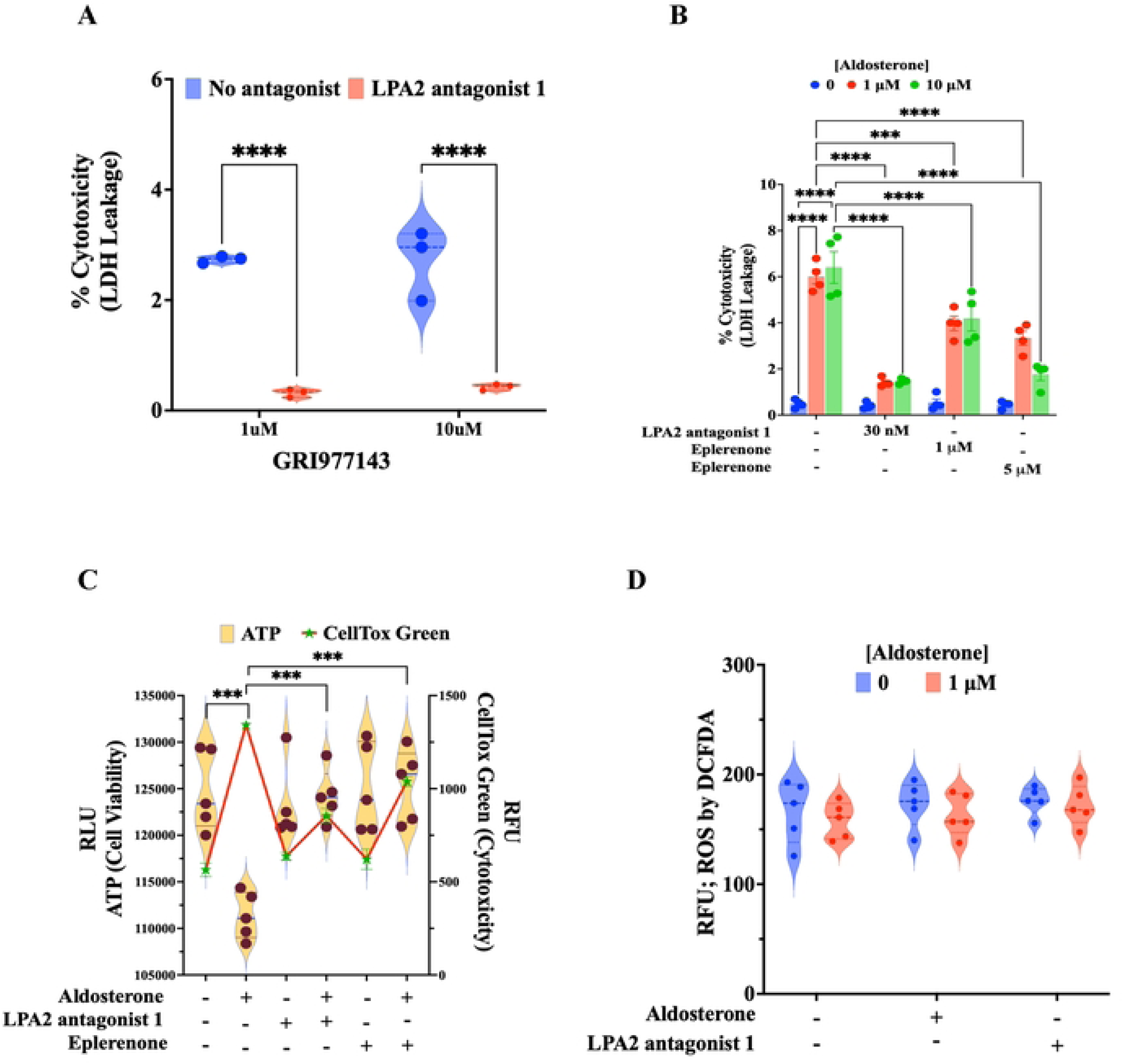
Aldosterone mediated cardiomyocyte cytotoxicity is MR and LPAR2 dependent in differentiated AC16 cells. Aldosterone (IμM) reduce cell viability, enhance cytotoxicity and induces plasma membrane damage as evidenced from LDH leakage was inhibited by either MR or LPAR2 antagonist in differentiated AC 16 human cardiomyocyte cells. Moreover, aldosterone did not show any changes to intracellular ROS accumulation in differentiated AC 16 cells. **A:** Effect of LPAR2 agonist GRI977143 on LDH leakage after 24 h, B: Effect of aldosterone on LDH leakage in presence or absence of MR and LPAR2 antagonist after 24 h, C: ATP cell viability and CellTox green cytotoxicity in differentiated AC 16 cells after 24 h aldosterone stimulation in presence or absence of MR and LPAR2 antagonist, D: Effect of aldosterone on intracellular ROS accumulation. Asterisks ***, **** indicates the significance level at *p* ≤0.001, *p* ≤ 0.0001, respectively.

We measured intracellular ATP content in the differentiated AC16 cells to further characterize cellular metabolic health and viability. Aldosterone treatment (24 h) markedly reduced intracellular ATP relative to controls, while pretreatment with either LPA2 antagonist 1 or eplerenone (30 min) prevented loss of ATP within the aldosterone treated cells. We used a CellTox Green cytotoxicity assay to confirm aldosterone elevates cell toxicity, and this was attenuated by either the LPAR2 or MR antagonists (**Fig. 5C**). Finally, given that aldosterone induces reactive oxygen species (ROS)-mediated cellular damage[46,47], we measured ROS levels using the DCFH-DA assay. However, no significant alterations in ROS generation were detected across any of the treatment conditions (**Fig. 5D**).

Collectively, these findings demonstrate that LPAR2 and MR work together in some fashion to mediate aldosterone-induced cardiomyocyte hypertrophy, cytotoxicity, pro-fibrotic collagen accumulation and metabolic toxicity. How the receptors mechanistically interact is not addressed in our current study.

## 4. Discussion

We identify lysophosphatidic acid receptor 2 (LPAR2) as a previously unrecognized GPCR signaling partner of aldosterone in cardiomyocytes. Although aldosterone is well-known to act through the canonical mineralocorticoid receptor[48–50], it is also thought to participate in rapid signaling through non-genomic signaling mediated by a plasma membrane receptor [15]. We find aldosterone activates GPCR LPAR2-mediated signaling to inhibit cAMP accumulation and to activate SRE. Aldosterone contributes to the damage of cardiomyocytes, in a manner that requires LPAR2 as well as the canonical MR. These findings provide a new framework to understand how aldosterone uses GPCR signaling along with classical MR signaling to affect cardiovascular remodeling, as occurs in cases of hyperaldosteronism and aging [15,51].

Previous work reports that aldosterone induces rapid activation of PI3K, PKC, ERK1/2, intracellular calcium, and cAMP signaling [15,20,52]. Although receptors such as GPER, AT1R, and EGFR transactivation are proposed to contribute to these rapid responses [19,31,52], there is no consensus on whether aldosterone directly participates in any membrane receptor activation [53,54]. Here we identified LPAR2 as a Gα_i/o_-coupled GPCR responsible for aldosterone-induced signaling in HEK293 cells based on a reporter of SRE activity, cAMP accumulation, pharmacological analyses, and a HEK293 line lacking Gi. The results with the ΔGi HEK293 cells are especially compelling because it shows Gα_i/o_ is essential for aldosterone to induce cAMP. Together these findings argue LPAR2 is as novel receptor participating in aldosterone-mediated signaling.

To put this novel aldosterone mediated signaling into a context of interorgan communication and pathology, we studied LPAR2 in AC16 human cardiomyocyte cells. We found the aldosterone mediated signaling in undifferentiated AC16 cells is only detectable after longer hormone exposure than seen in HEK293 cells. This delayed response in AC16 cells may reflect differences in receptor abundance, receptor trafficking, or a need to first induce GPCR accessory proteins. Notably, differentiated AC16 cells express 6-fold higher LPAR2 than proliferating AC16 cells, and they more rapidly responded to aldosterone. As well, over-expression of exogenous LPAR2 in undifferentiated AC16 cells accelerated the impacts of aldosterone on SRE and cAMP. The cell surface expression level of LPAR2 may be as a key factor for its aldosterone mediated response.

Activated GPCR are internalized in the process of receptor desensitization, recycling, and sustained intracellular signaling [44]. In AC16 cells, unlike HEK293 cells, aldosterone alone did not internalize LPAR2. However, aldosterone increased the amount of internalized receptor induced by the LPAR2 selective agonist GRI977143 [34]. Although aldosterone will also activate MR, we found that LPAR2 internalization induced by aldosterone plus GRI977143 could not be blocked by the MR antagonist spironolactone. Internalized LPAR2 is a marker of GPCR activation, and this is independent of MR. We speculate that aldosterone may be a positive allosteric modulator of LPAR2 that amplifies the activity induced by is its orthosteric agonist lysophosphatidic acid (modeled here by the selective GRI977143). Alternatively, aldosterone may impact the membrane environment about LPAR2 and thereby affect is interaction with LPA.

Our data also indicate there is a role for MR in the way aldosterone activates LPAR2. While aldosterone causes LPAR2 to internalize independent of MR, pharmacological inhibition of either LPAR2 or MR prevents aldosterone-induced cardiomyocyte injury, collagen, and hypertrophic gene expression. These observations suggest a two-phase signaling model: aldosterone rapidly activates LPAR2-dependent non-genomic signaling while it more slowly induces canonical MR mediated transcription. Both pathways are required for the pathological outcomes. In a serial model of these interactions, the rapid LPAR2 response to aldosterone is insufficient to induce fibrosis but could act to amplify the outputs of aldosterone-activated MR. Similar coordination has also been described for nuclear steroid receptors and GPCR pathways involving estrogen and glucocorticoid signaling [55,56].

Cardiac fibrosis is a pathological consequence of chronic excess aldosterone [9,57]. While pathology likely arises from the hemodynamic impacts of elevated aldosterone (REFS), this does not exclude the possibility of autonomous effects of the hormone upon cardiomyocytes. Although primary adult cardiomyocytes do not typically synthesize components of the fibrillar extracellular matrix[58,59], we observe basal Collagen 1 expression AC16 cells [46,47]. This expression may stem from the hybrid origin of the AC16 line, which was generated by fusing primary adult human ventricular cardiomyocytes with SV40-transformed human skin fibroblasts. Because fibroblasts are the primary producers of fibrillar collagens, AC16 cells retain genetic and epigenetic traits from their fibroblast parentage, permitting baseline or inducible Collagen 1 synthesis even following differentiation and SV40 T-antigen pathway silencing [32,60]. Importantly, we studied the impacts of aldosterone in the cardiomyocytes once the AC16 cells were differentiated, confirmed by robust expression of cardiac-specific markers (cTnI, *β*-MHC, and *α*-tubulin) alongside the absence of *α*-SMA [32,61–63]. Our data demonstrate that both aldosterone and direct LPAR2 activation increase collagen 1 protein expression, whereas pharmacological inhibition of either MR or LPAR2 abolishes aldosterone-induced collagen-1 accumulation as evident from ICC/IF. Lysophosphatidic acid signaling has previously been implicated in fibroblast activation, extracellular matrix remodeling, and tissue fibrosis in multiple organs, including the heart and kidney [29,64]. LPAR2 activates several downstream pathways, including RhoA/ROCK, MAPK, and PI3K/Akt signaling, all of which contribute to fibroblast activation and extracellular matrix synthesis [64]. Our findings suggest that aldosterone uses an existing pro-fibrotic LPAR2 signaling network within cardiomyocytes to amplify MR-driven fibrotic remodeling.

Independent of collagen 1, the hypertrophic response we observed in differentiated AC16 cells implies there is cooperative signaling between MR and LPAR2. Aldosterone exposure significantly increases MYH7 mRNA without affecting MYH6. Elevated MYH7/MYH6 is a widely accepted molecular indicator of pathological cardiac hypertrophy [45]. We find that inhibition of either LPAR2 or MR independently reduced this ratio in aldosterone treated AC16 cells; both rescued the hypertrophic phenotype. Interestingly, although aldosterone reduced cell viability based on intracellular ATP content, cytotoxicity, and elevated LDH leakage, we did not detect altered ROS. where aldosterone toxicity has traditionally been considered to increase oxidative stress [46]. Our observations suggest that early cardiomyocyte injury under these experimental conditions may occur independent of ROS.

Our study poses new problems to address in future work. Experiments were performed in cultured human cells, and we should like to validate the conclusions *in vivo* using genetic deletion or pharmacological inhibition of LPAR2 in mice. Pharmacological approaches were used to understand the relationship between aldosterone and LPAR2 involvement. A future step would generate biochemical data using ligand-binding assays, structural modeling, cryo-electron microscopy, or receptor mutagenesis to determine whether and how aldosterone directly interacts with LPAR2. Likewise, we infer aldosterone mediates an interaction between LPAR2 and MR. Work is needed to understand their interaction mechanisms, or how they together impact MR-dependent transcription.

## 5. Conclusion

This study identifies LPAR2 as a novel mediator of aldosterone-induced non-genomic signaling in a cardiomyocyte model and demonstrates that markers of pathological cardiac remodeling require coordinated activation of both LPAR2 and the classical MR pathway. Aldosterone activates membrane LPAR2-dependent Gαi/o signaling, leading to receptor internalization and rapid intracellular signaling and cooperates with MR to promote fibrosis, hypertrophy, and cardiomyocyte injury. We hypothesize that aldosterone first activates LPAR2, and this rapid signaling subsequently cooperates with MR-dependent transcription to cause cardiomyocyte injury. These findings expand the current understanding of aldosterone biology and identify a LPAR2-MR signaling axis as a potential therapeutic target for preventing aldosterone-induced myocardial injury.

## Credit authorship contribution statement

**Mohanty SK:** Conceptualization, methodology, investigation, formal analysis, writing original draft

**Tatar M:** Conceptualization, analysis, writing original draft, project administration, program funding

**Gong R:** Conceptualization, program funding

**Funding:** Funding for S.M, M.T. and R.G. was provided by National Institutes of Health, USA, U01AG086161

## Acknowledgement

We like to thank Prof. Asuka Inoue, from Graduate School of Pharmaceutical Science, Tohoku University, for providing ΔGi HEK cells.

## Data availability

The datasets generated and/or analyzed in this study are available from the corresponding author on reasonable request.

